# Extracellular ATP drives pancreatic cancer cell invasion via purinergic receptor-integrin interactions

**DOI:** 10.1101/2022.10.24.513477

**Authors:** Elena Tomas Bort, Megan D. Joseph, Qiaoying Wang, Edward P. Carter, Nicolas J. Roth, Jessica Gibson, Ariana Samadi, Hemant M. Kocher, Sabrina Simoncelli, Peter J. McCormick, Richard P. Grose

## Abstract

Pancreatic ductal adenocarcinoma (PDAC) is a cancer of unmet clinical need. Given the elevated ATP levels seen in PDAC, the purinergic axis represents an attractive therapeutic target. Mediated in part by highly druggable extracellular proteins, it plays essential roles in fibrosis, inflammation response and immune function. We have analysed the PDAC purinome using publicly available databases to discern which members may impact patient survival. We identified *P2RY2* to be the purinergic gene with the strongest association to hypoxia, the highest cancer cell specific expression and the strongest impact on overall survival. Invasion assays using a 3D spheroid model revealed P2Y_2_ to be critical in facilitating invasion driven by extracellular ATP. Using genetic modification or pharmacological strategies we identify the mechanism of this ATP-driven invasion to require direct protein-protein interactions between P2Y_2_ and αV integrins. Using DNA-PAINT super-resolution fluorescence microscopy, we found that P2Y_2_ regulates the amount and distribution of integrin αV in the plasma membrane. This work highlights a novel GPCR-integrin interaction in cancer invasion and its potential for therapeutic targeting.

## Introduction

Pancreatic ductal adenocarcinoma (PDAC), which accounts for 90% of diagnosed pancreatic cancer cases, has the lowest survival rate of all common solid malignancies. Surgery is the only potentially curative treatment, but unfortunately more than 80% of patients present with unresectable tumours (Kocher, 2022). Consequently, most patients survive less than 6 months after diagnosis, resulting in a 5-year survival rate of less than 5% when accounting for all disease stages (Bengtsson, Andersson and Ansari, 2020; Kocher, 2022). Despite continued efforts, this statistic has improved minimally in the past 50 years. Due to increasing incidence, late detection and lack of effective therapies, pancreatic cancer is predicted to be the second most common cause of cancer-related deaths by 2040 (Rahib *et al*., 2021).

Failure to improve clinical management significantly is mainly a result of chemoresistance (Neuzillet *et al*., 2017), thus it is of vital importance to find new therapeutics that can improve patient survival. PDAC is characterised by its desmoplastic stroma, with dense fibrosis leading to impaired vascularisation and high levels of hypoxia (Koong *et al*., 2000; Di Maggio *et al*., 2016). Lack of oxygen leads to cellular stress and death, resulting in the release of purines such as ATP and adenosine into the tumour microenvironment (Forrester and Williams, 1977; Pellegatti *et al*., 2008). PDAC has 200-fold more extracellular ATP than normal tissue (Hu *et al*., 2019), suggesting that purinergic signalling could represent an effective therapeutic target against PDAC.

The proteins underpinning purinergic signalling (the purinome), comprise several highly druggable membrane proteins involved in the regulation of extracellular purines, mainly ATP and adenosine (Burnstock and Novak, 2012; Boison and Yegutkin, 2019; Yu *et al*., 2021). Extracellular ATP is known to promote inflammation (Kurashima *et al*., 2012), growth (Ko *et al*., 2012) and cell movement (Martinez-Ramirez *et al*., 2016). Contrastingly, adenosine is anti-inflammatory and promotes immunosuppression (Schneider *et al*., 2021). There are ongoing clinical trials in several cancers, including PDAC, for drugs targeting the ectonucleotidase CD73 (NCT03454451, NCT03454451) and adenosine receptor 2A (NCT03454451) in combination with PD-1 checkpoint inhibitors and/or chemotherapy. However, a Phase II multi-cancer study evaluating an anti-CD73 and anti-PD-L1 combination was withdrawn due to minimal overall clinical activity (NCT04262388). This highlights the fact that further mechanistic understanding of purinergic signalling in PDAC is required to exploit its full therapeutic potential.

Here we combine bioinformatic, genetic and drug-based approaches to identify a novel mechanism mediating ATP-driven invasion, uncovering a new therapeutic target in PDAC. Beginning with an in-depth *in silico* analysis of the purinome of PDAC, using publicly available patient and cell line databases, we build on bioinformatic data associating the purinergic receptor P2Y_2_ with PDAC. After validating expression of P2Y_2_ in human PDAC cancers, we focused on identifying the function of the receptor in cancer cells. *In vitro* data underlined the importance of P2Y_2_ as a strong invasive driver, using a 3D physio-mimetic model of invasion. Finally, using a super-resolution imaging technique, DNA-PAINT, we characterise the behaviour of P2Y_2_ in the membrane at the single protein level, demonstrating the nanoscale distribution and interaction of this receptor with RGD-binding integrins in promoting pancreatic cancer invasion.

## Results

### The PDAC purinome associates with patient survival, hypoxia score and cell phenotype

The purinome encompasses 23 surface proteins, including pannexin 1, P2X ion channels, ectonucleotidases, and the P2Y and adenosine GPCRs (Di Virgilio *et al*., 2018) (Fig. 1A). Interrogating public databases, we determined which purinergic signalling genes significantly impact pancreatic cancer survival. First, we examined the pancreatic adenocarcinoma (PAAD) database from The Cancer Genome Atlas (TCGA; n=177 patients) analysing overall survival hazard ratios based on purinergic signalling gene expression (Fig. 1B). Expression of five purinergic genes correlated with decreased patient survival, with high *P2RY2* expression being associated with the highest hazard ratio (2.99, 95%, CI: 1.69 - 5.31, log-rank *p* = 8.5×10^-5^). We then examined the mutational profile and mRNA expression level of purinergic genes in patients. Using cBioPortal (Gao *et al*., 2013), we generated OncoPrints of purinergic signalling genes from PAAD TCGA samples (Sup. Fig. 1A), observing few genetic alterations in 0-3% of tumours and a heterogeneous percentage of tumours with high mRNA expression (z-score > 1) for each purinergic gene.

**Figure 1.**
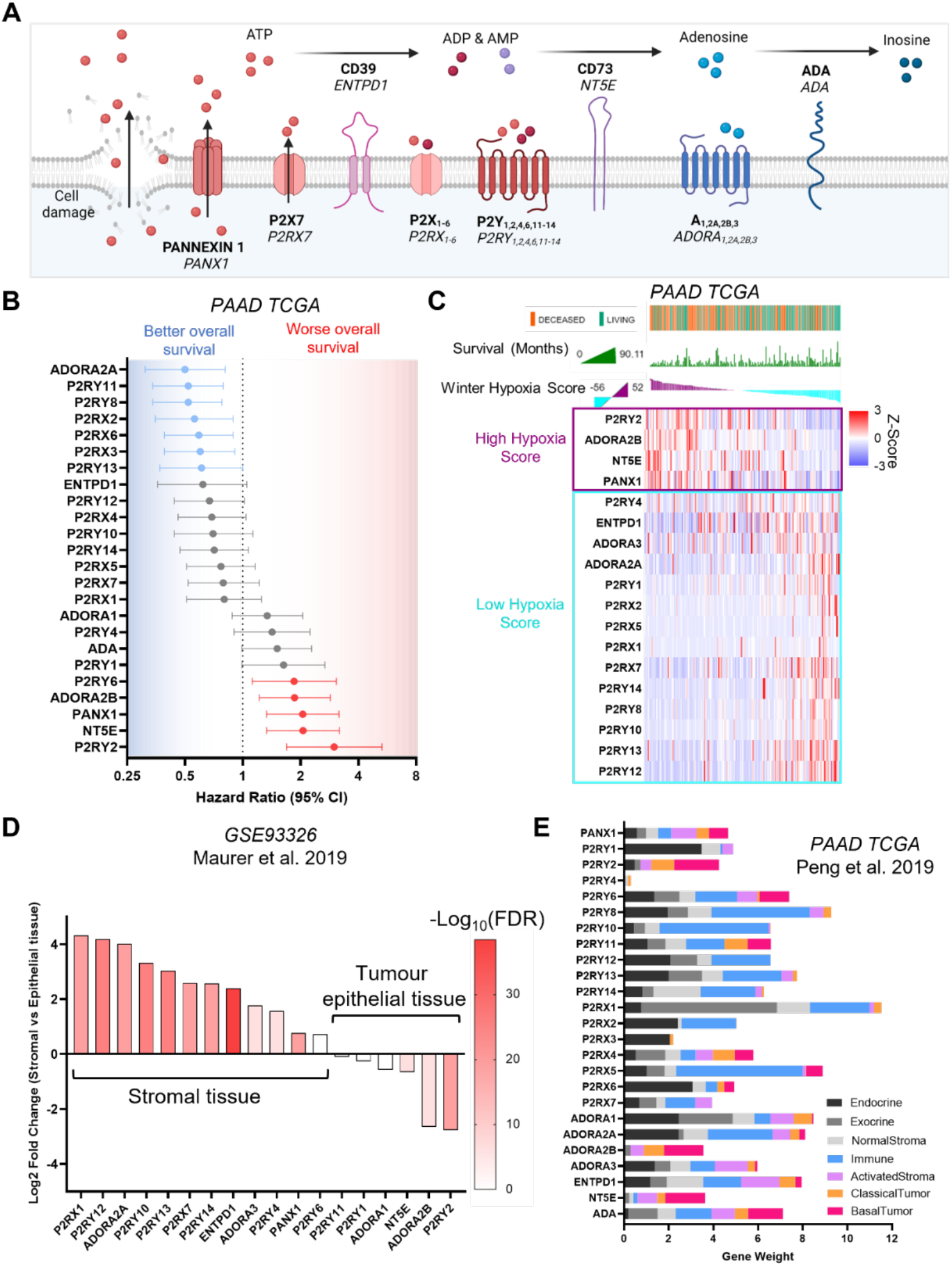
Characterisation of purinergic signalling in pancreatic adenocarcinoma. **A** Purinergic signalling proteins and gene names. **B** Hazard ratios of overall survival calculated using KMPlot and the PAAD TCGA cohort (n=177) for different purinergic genes. Statistically significant hazard ratios (log rank p-value) are highlighted in red for worse survival and in blue for better survival. **C** Heatmap of purinergic genes significantly correlated (*q* < 0.05) to high (purple) or low (light blue) Winter hypoxia scores in the PAAD TCGA data set. Overall survival status and overall survival in months is shown at the top, and samples are ranked using the Winter Hypoxia score (Generated with cBioPortal). **D** Differential expression analysis of 60 paired stromal and tumour tissue microdissections (GSE93326) showing significantly differentially expressed purinergic genes in stromal or tumour epithelial tissue. **E** Gene weights for purinergic genes representing the relevance of each gene to each cell type compartment, obtained from DECODER PDAC TCGA deconvolution analysis.

Since purinergic signalling has been strongly associated with hypoxia, the Winter (Winter *et al*., 2007), Ragnum (Ragnum *et al*., 2015) and Buffa (Buffa *et al*., 2010) hypoxia scores were used to examine the correlation between the expression of purinergic genes and HIF-1α in the PAAD TCGA database (Sup. Fig. 1B). Samples were divided into low (n=88) or high (n=89) hypoxia score, using the median hypoxia score to perform a differential expression analysis. CD73 (*NT5E*), adenosine A2B receptor (*ADORA2B*) and P2Y_2_ (*P2RY2*) mRNA expression associated strongly with the high hypoxia score group for all three hypoxia scores (log2 ratio >0.5, FDR <0.001). P2Y_2_ had the highest log2 ratio in all hypoxia signatures compared to other purinergic genes. With a more extensive gene signature, the Winter hypoxia score (99 genes) allowed for a more comprehensive relative hypoxia ranking of tumour samples, compared to Ragnum (32 genes) and Buffa (52 genes) signatures. Hence, we used cBioPortal to generate a transcriptomic heatmap of purinergic genes, ranked using the Winter hypoxia score and overlaid with overall survival data (Fig.1C). We observed a direct correlation between Winter hypoxia score and decreased overall survival for high hypoxia score-related purinergic genes.

We hypothesised that genes related to high hypoxia scores would be expressed preferentially in the tumour core. Mining published RNA-seq data from 60 paired PDAC samples of stroma and tumour microdissections (GSE93326) (Maurer *et al*., 2019) and performing differential expression analysis, we observed that most genes related to high Winter hypoxia scores (*P2RY2, ADORA2B* and *NT5E*) were expressed in the tumour epithelial tissue (Fig. 1D), except for *PANX1*, encoding for pannexin 1, which is involved in cellular ATP release (Bao, Locovei and Dahl, 2004).

To elucidate the cell type-specific purinergic expression landscape, we used published data from TCGA PAAD compartment deconvolution, using DECODER (Peng *et al*., 2019) to plot purinergic gene weights for each cell type compartment (Fig. 1E). The findings recapitulated the cell specificity data obtained from tumour microdissection analysis (Maurer *et al*., 2019) (Fig. 1D). Expression of purinergic genes in cancer cells was confirmed by plotting Z-scores of mRNA expression of PDAC cell lines from the cancer cell line encyclopaedia (Ghandi *et al*., 2019) (CCLE; Sup. Fig. 1D). Moreover, expression of purinergic genes in normal tissue from the Genotype-Tissue Expression (GTEx) database compared to cancer tissue (PAAD TCGA) also mimicked the results found with DECODER (Sup. Fig. 1D). *P2RY2*, encoding P2Y_2_ - a GPCR activated by ATP and UTP, was shown to be the purinergic gene most highly associated with cancer cell-specific expression in all our independent analyses (Fig. 1D, E; Sup. Fig. 1C, E). *P2RY2* additionally showed the strongest correlation with all hypoxia scores (Fig. 1C; Sup. Fig. 1B). Most importantly, of all purinergic genes, *P2RY2* expression had the biggest impact on adverse patient survival (Fig. 1B). These independent *in silico* analyses encouraged us to explore the influence of P2Y_2_ on pancreatic cancer cell behaviour.

### P2RY2 is expressed in cancer cells and causes cytoskeletal changes

To validate our bioinformatic findings, we performed RNAscope in human PDAC samples and corroborated P2Y_2_ mRNA expression as being restricted to the epithelial tumour cell compartment and not stroma, normal epithelium or endocrine tissues (n=3, representative images of 2 different patients shown in Fig. 2A and Sup. Fig. 2A). Using GEPIA (Tang *et al*., 2017), we analysed PAAD TCGA and GTEx mRNA expression of tumour (n=179) and normal samples (n=171). Tumour samples expressed significantly higher (*p* < 0.05) P2Y_2_ mRNA levels compared to the normal pancreas (Fig. 2B). Kaplan-Meier analysis from PAAD TCGA KMplot (Lánczky and Győrffy, 2021) showed a significant decrease in median overall survival in patients with high P2Y_2_ mRNA expression (median survival: 67.87 vs 17.27 months) (Fig. 2C).

**Figure 2.**
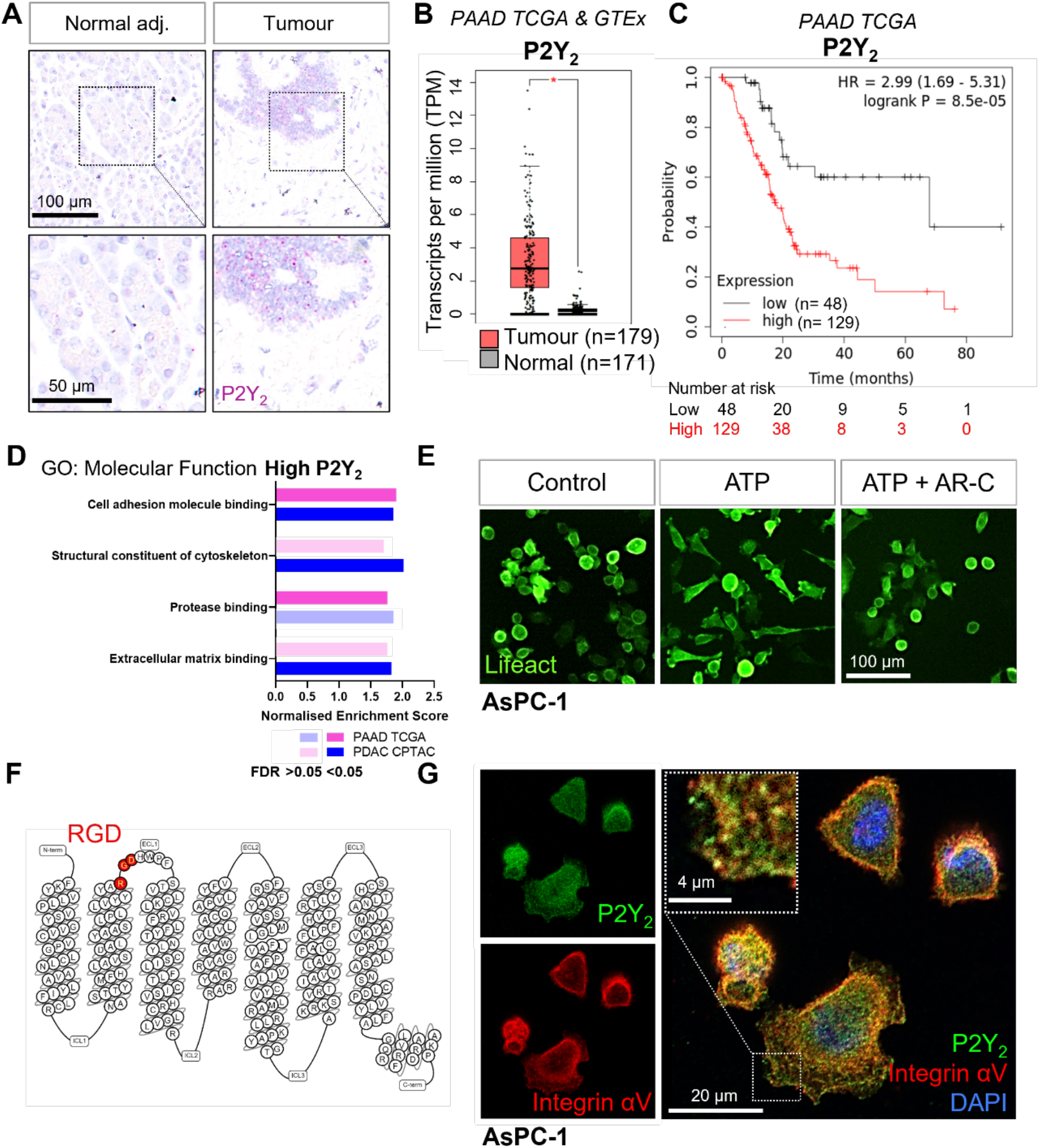
Expression of P2Y_2_ is specific to cancer cells, correlated with decreased overall survival in patients and drives cytoskeletal rearrangements. **A** RNAscope *in-situ* hybridisation of P2Y_2_ mRNA expression (magenta) in tumour and matching normal adjacent tissue. **B** P2Y_2_ mRNA expression in tumour (TCGA) and normal (GTEx) pancreatic tissue samples (* *p* <0.0001). Graph generated using GEPIA. **C** Kaplan-Meier plot comparing patients with high vs low expression of P2Y_2_ in the PAAD TCGA cohort. Graph generated using KMplot. **D** Top 4 results of a GSEA (performed with WebGestalt) of two different pancreatic adenocarcinoma patient cohorts (PAAD TCGA and PDAC CPTAC) for the ‘Molecular Function’ Gene Ontology (GO) functional database. **E** Incucyte images of the pancreatic cancer cell line AsPC-1 12 hours after treatment with 100 μM ATP alone or with 5 μM AR-C (P2Y2 antagonist). Cells are transduced with Lifeact to visualise f-actin (green). **F** Schematic of the amino acid sequence of P2Y_2_ showing an RGD motif in the first extracellular loop (image generated in gpcrdb.org). **G** IF staining of P2Y_2_ (green), integrin αV (red) and DAPI (blue) in AsPC-1 cells showing colocalization of P2Y_2_ and integrin αV (yellow).

To predict P2Y_2_ function in PDAC, we performed gene set enrichment analysis (GSEA) of high vs low mRNA expressing P2Y_2_ tumour samples, divided by the median expression, for PAAD TCGA (n=177) and the PDAC Clinical Proteomic Tumour Analysis Consortium (CPTAC) (n=140) databases. The top four enriched gene sets from the Gene Ontology ‘Molecular function’ functional database were associated with cell adhesion molecule binding, the cytoskeleton, protease binding and extracellular matrix binding (Fig. 2D). To initially evaluate the validity of the GSEA results *in vitro*, we used the PDAC cell line AsPC-1, transduced with Lifeact, which fluorescently labels filamentous actin structures (Riedl *et al*., 2008), and monitored cell morphology using the Incucyte live-cell analysis system. Cells treated with ATP (100 μM) showed cytoskeletal rearrangements which were blocked by the selective P2Y_2_ antagonist AR-C118925XX (AR-C; 5 μM; Fig 2E)(Muoboghare, Drummond and Kennedy, 2019).

P2Y_2_ is the only P2Y GPCR possessing an RGD motif, found in its first extracellular loop (Fig. 2F). Through this RGD motif, P2Y_2_ has been shown to interact with αV integrins (Erb *et al*., 2001), but the significance of this interaction has not been explored in cancer. Immunofluorescence (IF) showed colocalization of integrin αV and P2Y_2_ in the PDAC cell lines AsPC-1 and BxPC-3, while MIA PaCa-2 cells showed low expression of both proteins, and PANC-1 showed high integrin αV and low P2Y_2_, matching CCLE data (Fig 2G; Sup. Fig. 2B, C). We hypothesized that P2Y_2_, through its RGD motif, could engage αV integrins in cancer cells in the presence of ATP, leading to increased migration and invasion.

### Targeting P2Y_2_ and its RGD motif decreases ATP-driven invasion in PDAC cell lines

To evaluate the impact of P2Y_2_ in pancreatic cancer cell invasion, we used a 3D hanging drop spheroid model (Murray *et al*., 2022). PDAC cell lines were combined in a ratio of 1:2 (Kadaba *et al*., 2013), using an immortalised stellate cell line, PS-1 (Froeling *et al*., 2009) to form spheres (Fig. 3A), recapitulating the ratios of the two biggest cellular components in PDAC. Stellate cells also aid in sphere formation and cancer cell invasion (Murray *et al*., 2022). Spheres were embedded in a Collagen type I and Matrigel mix and cultured for 48 hours until imaging and fixing (Fig. 3A). Extracellular ATP concentration in tumours is in the hundred micromolar range (Pellegatti *et al*., 2008). Treating spheres with P2Y_2_ agonists ATP and UTP (100 μM) increased invasion of the PDAC cell line AsPC-1 significantly compared to vehicle control (*p* < 0.0001 and *p =* 0.0013 respectively), and this was blocked by the P2Y_2_ selective antagonist AR-C (5 μM, *p* = 0.0237 and *p* = 0.0133; Fig 3B, C; Sup. Fig. 3A). Importantly, a non-hydrolysable ATP (ATPγS;100 μM) showed similar effects, implicating ATP and not its metabolites as the cause of the invasion (Sup. Fig. 3B). Of note, IF staining of PS-1 cells showed negligible expression of P2Y_2_ (Sup. Fig. 3C). To determine whether integrin association was necessary for ATP-driven invasion, we treated spheres with 10 μM cyclic RGDfV peptide (cRGDfV), which binds predominantly to αVβ3 to block integrin binding to RGD motifs (Kapp *et al*., 2017), such as that in P2Y_2_ (Ibuka *et al*., 2015). cRGDfV treatment reduced ATP-driven motility significantly, both in 3D spheroid invasion assays (*p* < 0.001) (Fig 3B, C) and in 2D Incucyte migration assays (Sup. Fig. 3D, E). Treatment with AR-C decreased motility. To ensure that this behaviour was not restricted to AsPC-1 cells, experiments were corroborated in the BxPC-3 cell line (Sup. Fig. 3F, G).

**Figure 3.**
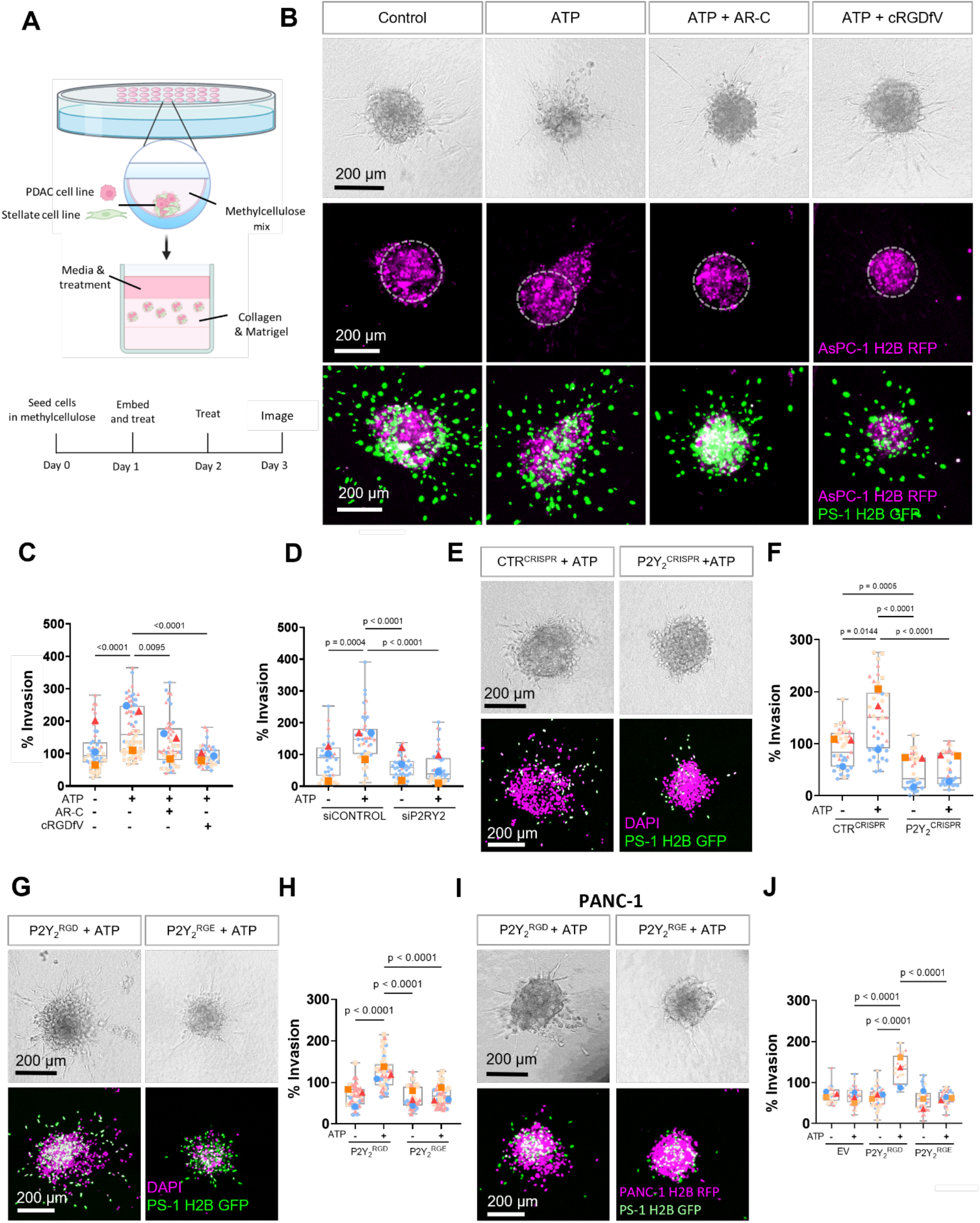
The RGD motif in P2Y_2_ is required for extracellular ATP-driven cancer cell invasion. **A** Schematic diagram of the hanging drop sphere model for 3D sphere invasion assays. **B** Brightfield and fluorescent images of spheres formed using AsPC-1 cells (magenta) with a histone 2B (H2B) tagged with red fluorescent protein (RFP) and the stellate cell line PS-1 (green) with H2B tagged with a green fluorescent protein (GFP). Middle pannel shows AsPC-1 cells in spheres with a dotted line highlighting the central sphere area. Spheres were treated with vehicle control or 100 μM ATP alone or with 5 μM AR-C or 10 μM cRGDfV. The quantification is shown in **C** using SuperPlots, where each colour represents a repeat and the larger points represent the mean % Invasion for each repeat. **D** Quantification of spheres formed by AsPC-1 cells transfected with a control siRNA or P2Y_2_ siRNA and treated with or without 100 μM ATP. **E** Brightfield and fluorescent images of spheres formed by AsPC-1 cells subjected to CRISPR/Cas9 gene disruption using a control guide RNA (CTR^CRISPR^) or P2Y_2_ guide RNAs (P2Y_2_^CRISPR^) and treated with or without 100 μM ATP. Quantification in **F**. **G, I** Brightfield and fluorescent images of AsPC-1 P2Y_2_^CRISPR^ cells or PANC-1 cells (respectively) transfected with wild-type *P2RY2* (P2Y_2_^RGD^) or mutant *P2RY2^D97E^* (P2Y_2_^RGE^) treated with or without 100 μM ATP and its quantification in **H** and **J**, respectively. Statistical analysis with Kuskal-Wallis multiple comparison test.

To further verify that ATP-driven invasion was dependent on P2Y_2_, we silenced P2Y_2_ expression in AsPC-1 cells using siRNA (Fig. 3D; Sup. Fig. 3H), abrogating the invasive response to ATP (*p* < 0.0001). P2Y_2_ involvement in this phenomenon was confirmed by generating a P2Y_2_ CRISPR-Cas9 AsPC-1 cell line (P2Y_2_^CRISPR^), which displayed a significant decrease in invasion compared to a control guide RNA CRISPR cell line (CTR^CRISPR^) in both ATP-treated (*p* = 0.0005) and non-treated (*p* < 0.001) conditions (Fig. 3F, E). These findings demonstrate that P2Y_2_ is essential for ATP-driven cancer cell invasion.

To determine the importance of the RGD motif of P2Y_2_ in ATP-driven invasion, we obtained a mutant P2Y_2_^D97E^ construct, where the RGD motif was replaced by RGE (P2Y_2_^RGE^), which has less affinity for αV integrins (Erb *et al*., 2001). This mutant was transfected into AsPC-1 P2Y_2_^CRISPR^ cells and compared to cells transfected with wild-type P2Y_2_ (P2Y_2_^RGD^; Sup. Fig. 3I). Only spheres containing P2Y_2_^RGD^ transfected cells demonstrated a rescue of the ATP-driven invasive phenotype (*p* < 0.0001; Fig. 3G, H). P2Y_2_^RGE^ spheres did not respond to ATP treatment. To ensure this behaviour was not influenced by CRISPR effects, we repeated the experiment with the PANC-1 cell line, which expressed very low levels of P2Y_2_, but high levels of integrin αV (Sup. Fig. 2B, C). No ATP-driven invasion was observed in PANC-1 cells transfected with an empty vector (EV) or with P2Y_2_^RGE^ (Fig. 3I,J). Only when transfecting PANC-1 cells with P2Y_2_^RGD^ was ATP-driven invasion observed (*p* < 0.001). These results demonstrate that the RGD motif of P2Y_2_ is required for ATP-driven cancer cell invasion.

### DNA-PAINT reveals RGD-dependent changes in P2Y_2_ and integrin αV surface expression

To interrogate how P2Y_2_ interacts with αV integrins, we examined the nanoscale organisation of P2Y_2_ and αV proteins under different treatment conditions using a multi-colour quantitative super-resolution fluorescence imaging method, DNA-PAINT. DNA-PAINT is a single-molecule localisation microscopy (SMLM) method based on the transient binding between two short single-stranded DNAs - the ‘imager’ and ‘docking’ strands. The imager strand is fluorescently labelled and freely diffusing in solution, whilst the docking strand is chemically coupled to antibodies targeting the protein of interest. For DNA-PAINT imaging of P2Y_2_ and integrin αV, proteins were labelled with primary antibodies chemically coupled to orthogonal docking sequences featuring a repetitive (ACC)n or (TCC)n motif, respectively (Fig. 4A). The benefit of such sequences is to increase the frequency of binding events, which in turn allows the use of relatively low imager strand concentrations without compromising overall imaging times, whilst achieving high signal-to-noise ratio and single-molecule localisation precision (Strauss and Jungmann, 2020).

**4.**
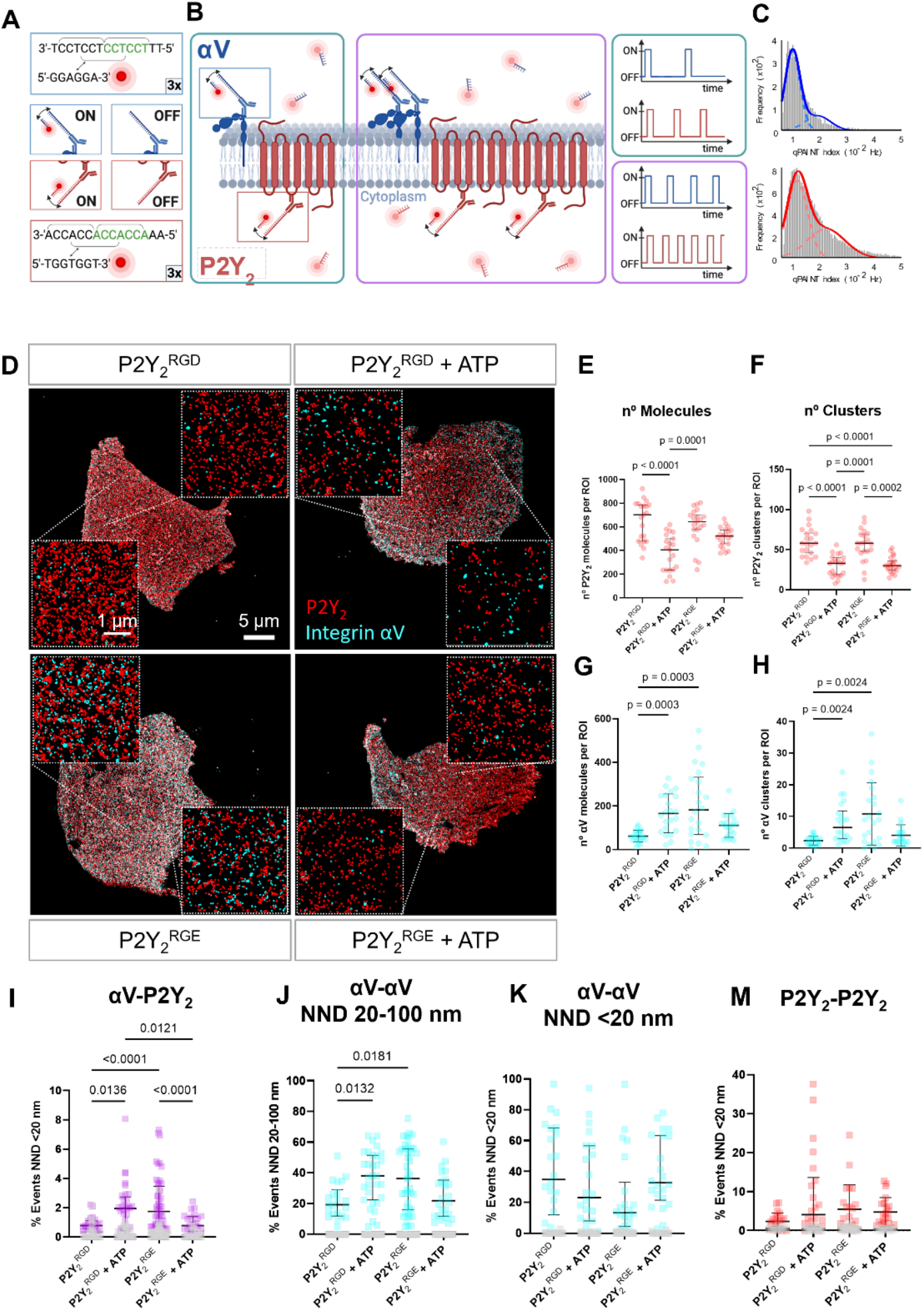
DNA-PAINT super-resolution microscopy reveals ATP and RGD-dependent changes in number and distribution of integrin αV and P2Y_2_ molecules in the plasma membrane. **A, B** Overview of the DNA-PAINT microscopy technique and qPAINT analysis pipeline**. C** Histogram of qPAINT indices for αV (blue) and P2Y_2_ (red) single molecule localisation clusters. Solid lines represent multi-peak Gaussian fit. **D** Rendered DNA-PAINT images of AsPC-1 P2Y_2_^CRISPR^ cells transfected with P2Y_2_^RGD^ or P2Y_2_^RGE^ with or without 100 μM of ATP and close ups showing the protein maps reconstructed from DNA-PAINT localization maps of P2Y_2_ (red) and integrin αV (cyan). The quantification of the number of proteins or protein clusters (>3 proteins) in each region of interest (ROI) are for P2Y_2_ (red)(**C** and **D** respectively) and integrin αV (cyan) (**E** and **F** respectively). Quantification of protein proximity using the nearest neighbour distance (NND), with the percentages of integrin αV and P2Y_2_ proteins being < 20 nm apart (**G**), between different αV integrins being 20-100 nm (**H**) or < 20 nm (**I**) apart; and P2Y_2_ from other P2Y_2_ proteins being < 20 nm apart (**J**). Statistical analysis with Kuskal-Wallis multiple comparison test.

The repetitive binding of imager and docking DNA strands in DNA-PAINT causes the same protein to be detected multiple times with nearly identical coordinates, originating a cluster of single molecule localisation around the true position of the protein. In contrast to other SMLM approaches, it is possible to take advantage of the DNA-binding kinetics to stoichiometrically calculate the number of proteins detected in each cluster of single molecule localisations, via an approach known as qPAINT (Schnitzbauer *et al*., 2017). As exemplified in Fig. 4B (and detailed in the methods section), qPAINT relies on the first order binding kinetics between individual imager and docking strands to determine the number copies of a protein that reside within a cluster of single-molecule localisations. The qPAINT index histograms obtained from P2Y_2_ and αV DNA-PAINT data sets were fitted with a multi-peak Gaussian function, identifying peaks located at multiples of a qPAINT index value of *q*_*i*,1_ 0.011 Hz and 0.009 for the P2Y_2_ and αV docking-imager pairs, respectively (Fig. 4C). These values were thus used to quantify the exact number of P2Y_2_ and αV proteins in all the clusters of single-molecule localisation in the DNA-PAINT data sets. By combining qPAINT with spatial statistics, we recovered a good estimation of the ground truth position of all the proteins in the DNA-PAINT data and quantified protein clustering.

We have previously analysed GPCR oligomerisation quantitively using DNA-PAINT super-resolution microscopy of P2Y_2_ in AsPC-1 cells (Joseph *et al*., 2021), where we observed a decrease in P2Y_2_ oligomerisation upon AR-C treatment. Hence, we questioned whether the RGD motif in P2Y_2_ affected receptor distribution and clustering. We imaged AsPC-1 P2Y_2_CRISPR cells transfected with P2Y_2_^RGD^ or P2Y_2_^RGE^ in the absence or presence of 100 μM ATP for 1 hour (Fig. 4D). We observed a 42% decrease in the median density of P2Y_2_ proteins at the membrane when P2Y_2_^RGD^ cells were treated with ATP, compared to control (*p* < 0.0001; Fig. 4E). In contrast, although a slight decrease in the density of P2Y_2_ proteins on P2Y_2_^RGE^ cells was observed following ATP treatment, it did not reach statistical significance (*p* = 0.1570). The density of P2Y_2_ proteins and protein clusters in both P2Y_2_^RGD^ and P2Y_2_^RGE^ controls were equivalent (Fig 4E, F; *p* > 0.9999), indicating similar expression of the receptor at the surface in both control conditions. Interestingly, the density of P2Y_2_ clusters decreased significantly in both conditions when treating with ATP (Fig. 4G; 43% decrease, *p* < 0.0001 for P2Y_2_^RGD^ and 48% decrease, *p* = 0.0002 for P2Y_2_^RGE^). We repeated these studies with AsPC-1 cells treated with ATP +/- cRGDfV, only observing a reduction of P2Y_2_ at the membrane with ATP alone (68% decrease, *p* <0.0001), while co-treatment with cRGDfV prevented this change (*p* > 0.9999; Sup. Fig. 4C, D). These findings highlight that the RGD motif is required for αV integrin to control P2Y_2_ levels at the membrane.

Moreover, we observed an increase in the density of αV molecules and αV molecules at the membrane when stimulating P2Y_2_^RGD^ with ATP (165 αV molecules/ROI, IQR = 162.75; 6.5 αV clusters/ROI, IQR = 8.75) compared to P2Y_2_^RGD^ without stimulation (58 αV molecules/ROI, IQR = 41; 2.5 αV clusters/ROI, IQR = 2; *p* = 0.0003; Fig. 4G, H). In absence of stimulation, P2Y_2_^RGE^ transfected cells exhibited more αV molecules and clusters at the membrane (182 αV molecules/ROI, IQR = 262.75; 9 αV clusters/ROI IQR = 14) compared to P2Y_2_^RGD^ cells (*p* = 0.0003, *p*=0.0024, respectively). However, treating P2Y_2_^RGE^ cells with ATP did not result in significant changes in αV molecules and clusters (112.5 αV molecules/ROI, IQR = 78.75; 3 clusters/ROI, IQR = 3). When the number of clusters was normalised with the number of αV molecules, to obtain the percentage of αV in clusters (Sup. Fig. 4A), there was no significant difference between conditions (*p* > 0.9999), indicating that the increase in the number of αV clusters was due to an increase in the number of αV proteins at the membrane. Taken together, these data indicate an RGD motif-dependent function of activated P2Y_2_ in localising integrin αV to the membrane.

By analysing nearest neighbour distance (NND) between proteins, we detected a higher percentage of integrin αV proteins in <20 nm proximity to P2Y_2_ in P2Y_2_^RGD^ cells following ATP stimulation (Fig. 4I; 62 % increase, *p* < 0.0001). In contrast, P2Y_2_^RGE^ cells stimulated with ATP showed a 71% decrease (*p* < 0.0001) in αV molecules in close proximity to P2Y_2_ in comparison to unstimulated cells. Analysing the percentage of αV proteins with NND in the range of 20-100 nm, we saw a similar pattern (Fig. 4J). ATP-stimulated P2Y_2_^RGD^ and unstimulated P2Y_2_^RGE^ cells showed an increased percentage of αV proteins spaced at this range compared to untreated P2Y_2_^RGD^ cells (136% increase with *p* = 0.0024 and 177% increase with *p* = 0.0004). No significant changes were observed in NND of <20 nm between αV proteins in any of the conditions (Fig. 4K), and the same was true for P2Y_2_ with respect to other P2Y_2_ proteins (Fig. 4M). In summary, our SMLM studies demonstrate a reciprocal interaction between αV integrin and P2Y_2_ receptors, where P2Y_2_ can alter integrin localisation to the plasma membrane while αV integrins influence activated P2Y_2_ internalisation.

## Discussion

Improved biological mechanistic understanding of PDAC is vital to identify effective therapeutic approaches to improve patient survival. Purinergic signalling includes many druggable targets that have been related to hypoxia (Synnestvedt *et al*., 2002), immunosuppression (Fong *et al*., 2020), and invasion (Li *et al*., 2015), but have been relatively underexplored in PDAC. In this study, we used publicly available databases to identify purinergic signalling genes that could be promising targets for PDAC, determining P2Y_2_ as a driver of pancreatic cancer cell invasion. Extracellular ATP increased invasion in a 3D spheroid model of PDAC, which was blocked by targeting P2Y_2_ genetically and pharmacologically. Mechanistically, we identified that the RGD motif in the first extracellular loop of P2Y_2_ is required for ATP-driven cancer invasion. Importantly, quantitative DNA-PAINT super-resolution fluorescence microscopy revealed a role of this RGD motif in orchestrating the number of P2Y_2_ and αV integrin proteins at the plasma the membrane, upon ATP stimulation.

Purinergic signalling has been associated classically with hypoxia and immune function in cancer (Di Virgilio *et al*., 2018). While expression of most purinergic genes was associated predominantly with immune cells and low hypoxia scores (Fig. 1C, E), expression of genes correlated with worse survival and hypoxia (*NT5E, ADORA2B* and *P2RY2*) was associated with tumour cells. The role of CD73 in PDAC has been examined in several studies (Yu *et al*., 2021) (NCT03454451, NCT03454451). In contrast, adenosine A2B receptor has not been well studied and this is the first report of its correlation with decreased overall survival in PDAC. From our analyses, P2Y_2_ was associated with the worst patient overall survival, highest patient hypoxia scores and strongest correlation to cancer cell expression. These observations were supported by published immunohistochemical staining showing that P2Y_2_ localised predominantly in cancer cells in human PDAC and that P2Y_2_ activation led to HIF-1α expression (Hu *et al*., 2019). Hence, we decided here to explore P2Y_2_ in greater depth.

P2Y_2_ has been associated with cancer cell growth and glycolysis in PDAC (Ko *et al*., 2012; Hu *et al*., 2019; Wang *et al*., 2020). Surprisingly, our GSEA results in two different cohorts suggested a possible additional function in invasion. Increased glycolysis and cytoskeletal rearrangements have been linked (Park *et al*., 2020), and both events could occur downstream of P2Y_2_ activation. P2Y_2_ has been implicated in invasive phenotypes in prostate, breast and ovarian cancer (Jin *et al*., 2014; Li *et al*., 2015; Martinez-Ramirez *et al*., 2016). Moreover, many genes sets associated with P2Y_2_ expression were related to integrin signalling. The RGD motif in the first extracellular loop of P2Y_2_ results in a direct interaction of P2Y_2_ with RGD-binding integrins, particularly integrins αVβ3 and αVβ5 (Erb *et al*., 2001). This can exert phenotypic effects – for example, binding of P2Y_2_ to integrins via its RGD motif is necessary for tubule formation in epithelial intestinal cell line 3D models (Ibuka *et al*., 2015). We focus here on the importance of the RGD motif of P2Y_2_ in a cancer context. Treating 3D PDAC spheroids with ATP resulted in increased invasion (Fig. 3), which was blocked by the selective P2Y_2_ antagonist AR-C and by the selective αVβ3 cyclic RGD-mimetic peptide inhibitor cRGDfV. Likewise, spheres containing ASPC-1 P2Y_2_^CRISPR^ or PANC-1 cells transfected with mutant P2Y_2_^RGE^, which decreases the affinity of P2Y_2_ for integrins, did not show increased invasion upon ATP stimulation. Altogether, these results support P2Y_2_ involvement in PDAC cell invasion and show the RGD motif is essential for this function. Despite efforts, there are currently no clinically efficacious P2Y_2_ antagonists, with poor oral bioavailability and low selectivity being major issues (Neumann *et al*., 2022). Our findings demonstrate that P2Y_2_ can also be targeted by blocking its interaction with RGD-binding integrins, due to its dependence on integrins for its pro-invasive function.

GPCR-integrin crosstalk is involved in many biological processes (Wang *et al*., 2005; Teoh *et al*., 2012). Only one study has directly examined the spatial distribution of integrins and GPCRs, however this relied on IF analysis (Erb *et al*., 2001), where only changes in the micron scale will be perceived, hence losing information of the nanoscale distances and individual protein interactions. In this work, we present a method to image integrin and GPCR dynamics using the recent quantitative DNA-PAINT super-resolution fluorescence microscopy technique (Schnitzbauer *et al*., 2017), which enabled us to look at and quantify the organisation of P2Y_2_ and integrin αV at the single protein level. We noted that upon ATP stimulation, the number of P2Y_2_ proteins at the plasma membrane decreased significantly after one hour, implying receptor internalisation. This phenomenon has been observed, were staining of P2Y_2_ at the membrane was reduced significantly after one hour of UTP stimulation (Tulapurkar *et al*., 2005). Of note, cytoskeletal rearrangements, which we have also observed upon ATP stimulation (Fig. 2E), were required for P2Y_2_ clathrin-mediated internalisation and authors noted that P2Y_2_ was most likely in a complex with integrins and extracellular matrix-binding proteins. Cells expressing RGE mutant P2Y_2_ or treated with cRGDfV, did not show significant changes in P2Y_2_ levels at the membrane upon ATP treatment. These results implicate the RGD motif in P2Y_2_ in agonist-dependent receptor internalisation.

P2Y_2_ affecting cell surface redistribution of αV integrin has been reported, with αV integrin clusters observed after 5 min stimulation with UTP (Chorna *et al*., 2007). We observed an increased number of αV integrin molecules and clusters one hour after ATP stimulation, although this increase in clusters was mainly due to the increase in total number of αV integrins at the membrane. The distance between αV integrin and P2Y_2_ decreased (NND < 20 nm) with ATP stimulation, indicating possible interaction. In contrast, with mutant P2Y_2_^RGE^, no significant changes in the number of P2Y_2_ or αV integrin proteins at the membrane were observed when stimulating with ATP. In fact, the number of αV integrins at the surface increased compared to wild-type. The same phenomenon was observed when treating normal AsPC-1 cells with cRGDfV and ATP. We speculate that the RGD in P2Y_2_ may regulate integrin-dependent P2Y_2_ internalisation and downstream signalling (Fig. 5).

**Figure 5.**
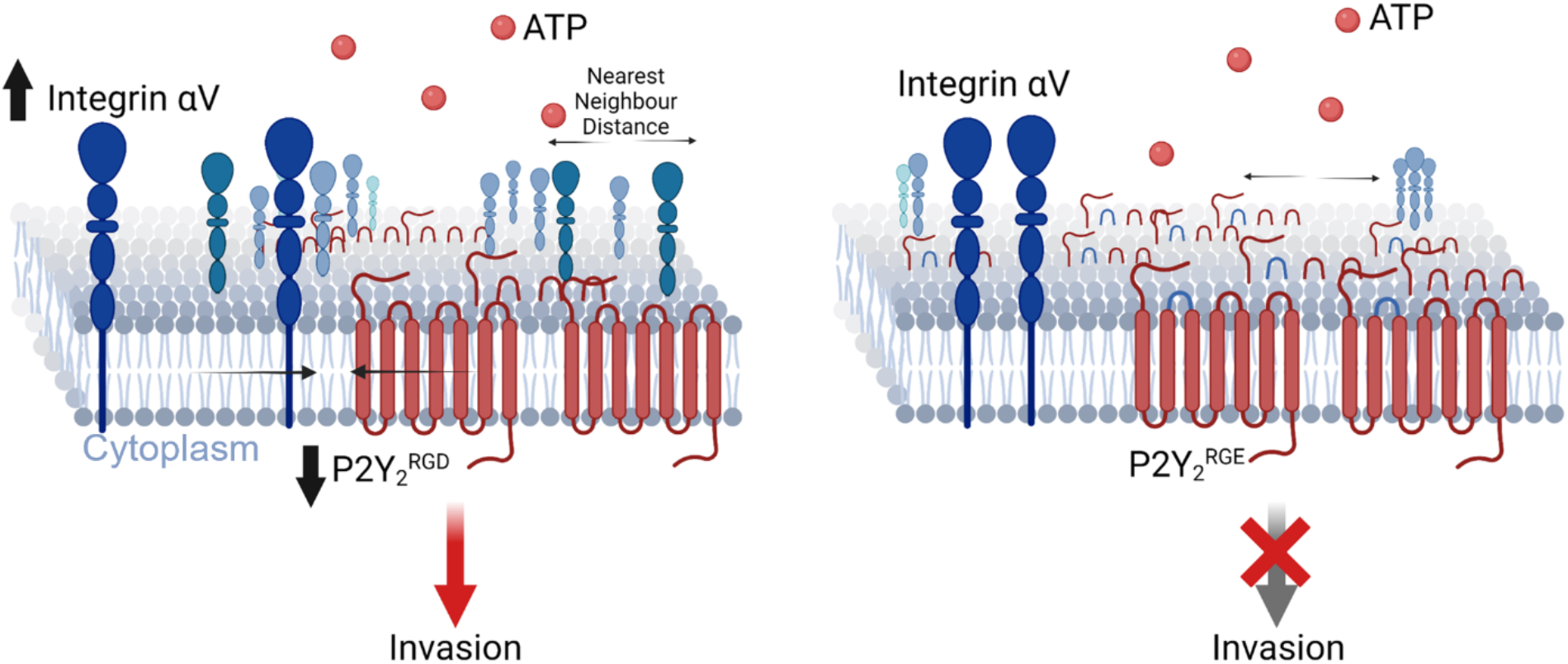
P2Y_2_ stimulation with extracellular ATP leads to RGD-dependent changes in number and distribution of integrin αV and P2Y_2_ and cancer cell invasion.

In summary, our study demonstrates that P2Y2, via its RGD motif, has a pivotal role in ATP-induced PDAC invasion through interacting with, and regulating the number of, αV integrins at the plasma membrane, making it a promising therapeutic target.

## Methods

### Data mining and bioinformatic analysis

Hazard ratios and the P2Y_2_ Kaplan-Meier plot for overall survival were obtained using Kaplan-Meier Plotter (Lánczky and Győrffy, 2021) and the pancreatic adenocarcinoma dataset from the cancer genome atlas (PAAD TCGA).

Using cBioPortal (Gao *et al*., 2013) and the database PAAD TCGA, mRNA differential expression analysis was performed for each Hypoxia Score (Buffa, Ragnum or Winter)(Winter *et al*., 2007; Buffa *et al*., 2010; Ragnum *et al*., 2015) by separating patients using the median hypoxia score. Results from purinergic genes were plotted in a volcano plot using VolcaNoseR (Goedhart and Luijsterburg, 2020). Significant hits were plotted in a heat map using cBioPortal. RNAseq raw counts from stromal and epithelial PDAC tissue from microdissections were downloaded from the GEO database (GSE93326)(Maurer *et al*., 2019) and a differential expression analysis was performed using DESeq2 (Love, Huber and Anders, 2014; Varet *et al*., 2016) in R.

Gene weight results from DECODER from PDAC tissues in the TCGA database were obtained from published results (Peng *et al*., 2019). Using GEPIA (Tang *et al*., 2017), mRNA expression of purinergic genes in normal tissue from the Genotype-Tissue Expression (GTEx) compared to cancer tissue (PAAD TCGA) were obtained. PDAC cell line mRNA z-scores or mRNA reads per kilobase million (RPKM) were obtained using cBioPortal and the Cancer Cell Line Encyclopaedia (CCLE) data.

For gene set enrichment analysis (GSEA), cBioPortal was used to separate PAAD TCGA or PDAC CPTAC patients into high and low *P2RY2* by *P2RY2* median expression and perform the differential expression analysis. Log ratio values were inserted in the WEB-based Gene SeT AnaLysis Toolkit (WebGestalt)(Liao *et al*., 2019), where ‘GO:Molecular Function’ with default analysis parameters selected.

### RNAscope® *in-situ* hybridisation

Formalin fixed paraffin embedded (FFPE) sections (n=3) of PDAC with stroma and normal adjacent tissue were obtained from the Barts Pancreas Tissue Bank (Project 2021/02/QM/RG/E/FFPE). These studies were approved by and performed in accordance with nationally required ethical standards (Hampshire B Research Ethics Committee: 18/SC/0630). Sections were stained using the human *P2RY2* probe (853761, ACD) and the RNAscope® 2.5 HD Assay-RED (ACD) following manufacturer’s instructions. Slides were imaged by NanoZoomer S210 slide scanner (Hamamatsu).

### Cell lines and cell culture

The pancreatic cancer cell lines AsPC-1, BxPC-3, MIA PaCa-2 and PANC-1 were kindly donated by Prof. Hemant Kocher (Queen Mary University of London), in addition to the immortalised stellate cell line PS-1 (Froeling *et al*., 2009). Cell lines expressing stable fluorescently labelled histone subunits (H2B) or Lifeact (Riedl *et al*., 2008) were transduced with viral supernatant obtained from HEK293T cells co-transfected with pCMVR8.2 (Addgene #12263) and pMD2.G (Addgene #12259) packaging plasmids, and either H2B-GFP (Addgene #11680), H2B-RFP (Addgene #26001) or Lifeact-EGFP (Addgene # 84383) plasmids using FuGENE transfection reagent (Promega), following manufacturer’s guidelines. Successfully transfected cells were isolated using a BD FACS Aria Fusion cell sorter. AsPC-1 P2Y_2_^CRISRP^ cells were generated by transfecting cells with a dual gRNA (TGAAGGGCCAGTGGTCGCCGCGG and CATCAGCGTGCACCGGTGTCTGG) CRISPR-CAS9 plasmid (VectorBuilder) with a mCherry marker which was used to select successfully transfected cells as above. Clonal expansion of single sorted cells was achieved with serial dilution cloning. Clones were evaluated by IF for P2Y_2_ compared to parental AsPC-1 cells. Cell lines were grown at 37 °C with 5% CO_2_ in DMEM (Gibco), RPMI-1640 (Gibco) or DMEM/F-12 (Sigma) supplemented with 10% fetal bovine serum (Sigma). Cells were monitored for mycoplasma contamination every six months.

### Cell fixation and immunofluorescent staining

Cells were seeded in coverslips placed in a 6 well-plate (Corning) and fixed the next day in 4% paraformaldehyde (LifeTech) for 30 min and washed 3x with phosphate buffered saline (PBS). Coverslips were placed in 0.1% Triton X-100 (Avantor) for 10 min for permeabilization, followed by 3 PBS washes and blocking with 5% bovine serum albumin (BSA; Merck) for 1 hour. Coverslips were incubated at 4 °C overnight with anti-P2Y2 (APR-010, Alomone labs) and anti-integrin αV antibodies (P2W7, Santa Cruz) diluted in blocking solution (1:100 and 1:200, respectively). After 3 PBS washes, coverslips were incubated for 1 hour with Alexa Fluor 647 goat anti-mouse and Alexa Fluor 488 goat anti-rabbit (Invitrogen) or Alexa Fluor 546 goat anti-rabbit at 1:1000, diluted in blocking buffer. Following 3 PBS washes, 4’,6-diamidino-2-phenylindole (DAPI, Sigma-Aldrich) was used as a nuclear stain and was incubated for 10 min. Slides were mounted using Mowiol (Calbiochem) and imaged 24 hours later using a LSM 710 confocal microscope (Zeiss).

### siRNA and plasmid transfection

Cells were seeded in 6 well plates at a density of 200,000 cells/well 24 hours before transfection. For siRNA experiments, cells were transfected with 20 nM pooled control or P2Y2-targeting siRNAs from a siGENOME SMARTpool (Dharmacon, GE Heathcare) with Lipofectamine 3000 (Invitrogen) following manufacturer’s instructions. For P2Y_2_ plasmid expression experiments, cells were transfected with 500 nM *P2RY2* (P2Y_2_^RGD^) or *P2RY2D97E* (P2Y_2_^RGE^) in pcDNA3.1 vector (Obtained from GenScript) or pcDNA3.1 alone (Empty vector, EV) together with lipofectamine 3000 and p3000 reagent (Invitrogen) as per manufacturer’s instructions. Plasmid concentration was selected by comparing AsPC-1 IF staining of P2Y_2_ with IF staining in AsPC-1 P2Y_2_^CRISPR^ and PANC-1 cells with different concentrations of plasmid to achieve a similar IF signal. Cells were split 48 hours post-transfection for experiments or imaged 72 hours post-transfection.

### 3D sphere model invasion assay

Spheres of PDAC cell lines with PS-1 cells were generated as described (Murray *et al*., 2022). Cancer cells at 22,000 cells/ml and PS-1 cells at 44,000 cells/ml were combined with DMEM/F-12 and 1.2% methylcellulose in a 4:1 ratio of methylcellulose (Sigma-Aldrich) and 20 μl drops, each containing 1000 cells, pipetted on the underside of a 15 cm dish lid (Corning) and hanging drops were incubated overnight at 37 °C. The next day, spheres were collected and centrifuged at 300 g for 4 minutes and washed with medium. A mix of 2 mg/ml collagen (Corning), 175 μl/ml Matrigel, 25 μl/ml HEPES (1M, pH 7.5) and 1N NaOH (for neutral pH correction) was prepared with DMEM/F12 medium. Spheroids were re-suspended and seeded in low attachment 96-well plates (50 μl per well) with 40 μl previously gelled mix in the bottom of the wells. Once set, 150 μl of DMEM/F12 was added with treatments. Spheres were treated with 100 μM adenosine 5’-triphosphate trisodium salt hydrate (ATP, Sigma), uridine 5’-triphosphate trisodium salt hydrate (UTP, Sigma) or adenosine 5’-[γ-thio]triphosphate tetralithium salt (ATPγS, Tocris) alone or with 5 μM AR-C118925XX (AR-C, Tocris) or 10 μM cycloRGDfV (cRGDfV, Sigma-Aldrich). Treatments were repeated 24 hours later. Spheres were imaged with a Zeiss Axiovert 135 light microscope at x10 on day 2 after seeding. Cells were stained with 4’,6-diamidino-2-fenilindol (DAPI) (1:1000) for 10 minutes and imaged with a Zeiss LSM 710 confocal microscope. %Invasion was calculated by drawing an outline around the total area *A_total_* and central area *A_central_* of the spheres with ImageJ (Fiji) and using the equation:

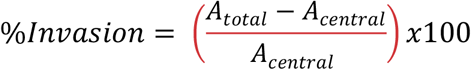

Results were plotted in SuperPlots by assigning different colours to repeats and superimposing a graph of average % Invasion with a darker shade of the assigned colour as described previously (Lord *et al*., 2020).

### Incucyte migration assay

In Incucyte ClearView 96-well cell migration plates (Essen BioScience), 40 μL medium with 5,000 cells were seeded in each well. A solution of 20 μL medium with 15 μM AR-C or 30 μM cRGDfV was added on top of the wells to achieve a final concentration of 5 μM and 10 μM respectively. Cells were allowed to settle for 15 minutes at room temperature and then placed at 37 °C for pre-incubation with the treatments for another 15 min. A volume of 200 μL of medium with or without 100 μM ATP was added in the appropriate reservoir wells and the plate was placed in the IncuCyte S3 (Essen BioScience) and was monitored every 4 hours for 39 hours (average doubling time of AsPC-1 cells (Chen *et al*., 1982)). Using the IncuCyte S3 2019A software, the migration index was calculated by analysing the average area occupied by the cells in the bottom well and was averaged with the initial average area occupied by cells in the top well.

### RNA extraction and qPCR analysis

RNA was extracted using the Monarch RNA extraction kit (BioLabs) as instructed by the manufacturer. The extracted RNA was quantified using a Nanodrop One Spectrophotometer (ThermoFisher Scientific). Using LunaScript RT Supermix kit (BioLabs), cDNA was prepared in a 20 μL reaction according to manufacturer’s instructions. The resulting cDNA was used in conjunction with MegaMix-Blue and *P2RY2* primers (Eurogentec; Forward sequence: GCTACAGGTGCCGCTTCAAC, reverse sequence: AGACACAGCCAGGTGGAACAT)(Hu *et al*., 2019) for quantitative polymerase chain reaction (qPCR) at the manufacturer’s recommended settings in a StepOnePlus Real-Time PCR System (Applied Biosystems). The relative mRNA expression was calculated using the 2^-ΔΔct^ method (Livak and Schmittgen, 2001) and normalised to GAPDH.

### DNA-antibody coupling reaction

DNA labelling of anti-aV antibody (P2W7, Santa Cruz) and anti-P2Y2 receptor antibody (APR-010, Alomone labs) was performed via maleimidePEG2-succinimidyl ester coupling reaction as previously described (Simoncelli *et al*., 2020; Joseph *et al*., 2021). Firstly, 30 μL of 250 mM DDT was added to 13 μL of 1 mM thiolated DNA sequences 5’-Thiol-AAACCACCACCACCA-3’ (Docking 1), and 5-Thiol-TTTCCTCCTCCTCCT-3’ (Docking 2) (Eurofins). The reduction reaction occurred under shaking conditions for 2 hours. 30 min after the reduction of the thiol-DNA started, 175 μL of 0.8 mg/mL antibody solutions were incubated with 0.9 μL of 23.5 mM maleimide-PEG2-succinimidyl ester cross-linker solution (Sigma-Aldrich) on a shaker for 90 min at 4 °C in the dark. Prior DNA-antibody conjugation, both sets of reactions were purified using Microspin Illustra G-25 columns (GE Healthcare) and Zeba spin desalting columns (7K MWCO, Thermo Fisher Scientific), respectively, to remove excess reactants. Next, coupling of anti-P2Y2 with with DNA docking 1 and anti-aV with DNA Docking 2 was performed by mixing the respective flow-through of the columns and incubate them overnight, in the dark, at 4°C under shaking. Excess DNA was removed via Amicon spin filtration (100K, Merck) and antibody-DNA concentration was measured using a NanoDrop One spectrophotometer (Thermo Fisher Scientific) and adjusted to 10 μM with PBS. Likewise, spectrophotometric analysis was performed to quantify the DNA-antibody coupling ratio and found to be ~1.2 in average for both the oligo-coupled primary antibodies.

### Cell fixation and immunofluorescence staining for DNA-PAINT imaging

Cells were seeded at 30,000 cells per channel on a six-channel glass bottomed microscopy chamber (μ-SlideVI^0.5^, Ibidi) pre-coated with rat tail collagen type I (Corning). The chamber was incubated at 37 °C for 8 hours before treatments. Cells were treated with 100 μM of ATP (or the equivalent volume of PBS as control) in medium for 1 hour and were fixed and permeabilised as described in the ‘Cell fixation and immunofluorescent staining’ section. Following permeabilisation, samples were treated with 50 mM ammonium chloride solution (Avantor) for 5–10 min to quench auto-fluorescence and cells were washed 3× in PBS. Blocking was completed via incubation with 5% BSA (Merck) solution for 1 hour followed by overnight incubation at 4°C with 1:100 dilutions of DNA labelled anti-P2Y2, and DNA labelled anti-αV antibody in blocking solution. The next day, samples were washed 3× in PBS and 150 nm gold nanoparticles (Sigma-Aldrich) were added for 15 min to act as fiducial markers for drift correction, excess of nanoparticles was removed by 3× washes with PBS. Samples were then left in DNA-PAINT imager buffer solution, prepared as described below, and immediately used for DNA-PAINT imaging experiments.

### DNA-PAINT imager solutions

A 0.1 nM P2Y_2_ imager strand buffer solution (5-TTGTGGT-3’-Atto643, Eurofins) and a 0.2 nM αV imager strand buffer solution (5-GGAGGA-3’-Atto643, Eurofins) were made using 1× PCA (Sigma-Aldrich), 1× PCD (Sigma-Aldrich), 1× Trolox (Sigma-Aldrich), 1× PBS and 500 mM NaCl (Merck) which facilitates establishment of an oxygen scavenging and triplet state quencher system. Solutions were incubated for 1 h in the dark before use. Stock solutions of PCA, PCD and Trolox were prepared as follows: 40× PCA (protocatechuic acid) stock was made from 154 mg of PCA (Sigma-Aldrich) in 10 mL of Ultrapure Distilled water (Invitrogen) adjusted to pH 9.0 with NaOH (Avantor, Radnor Township, PA, USA). 100x PCD (protocatechuate 3,4-dioxygenase) solution was made by adding 2.2 mg of PCD (Sigma-Aldrich) to 3.4 mL of 50% glycerol (Sigma-Aldrich) with 50 mM KCl (Sigma-Aldrich), 1 mM EDTA (Invitrogen), and 100 mM Tris buffer (Avantor). 100x Trolox solution was made by dissolving 100 mg of Trolox (Sigma-Aldrich) in 0.43 mL methanol (Sigma-Aldrich), 0.345 mL 1 M NaOH, and 3.2 mL of Ultrapure Distilled water (Invitrogen, Waltham).

### Exchange-PAINT Imaging Experiments

Exchange DNA-PAINT imaging was performed on a custom built total internal reflection fluorescence (TIRF) microscope based on a Nikon Eclipse Ti-2 microscope (Nikon Instruments) equipped with a 100× oil immersion TIRF objective (Apo TIRF, NA 1.49) and a Perfect Focus System. Samples were imaged under flat-top TIRF illumination with a 647 nm laser (Coherent OBIS LX, 120 mW), that was magnified with custom-built telescopes, before passing through a beam shaper device (piShaper 6_6_VIS, AdlOptica) to transform the Gaussian profile of the beam into a collimated flat-top profile. The beam was focused into the back focal plane of the microscope objective using a suitable lens (AC508-300-A-ML, Thorlabs), passed through a clean-up filter (FF01-390/482/563/640-25, Semrock) and coupled into the objective using a beam splitter (Di03-R405/488/561/635-t1-25×36, Semrock). Laser polarization was adjusted to circular after the objective. Fluorescence light was spectrally filtered with an emission filter (FF01-446/523/600/677-25, Semrock) and imaged on a sCMOS camera (ORCA-Flash4.0 V3 Digital, Hamamatsu) without further magnification, resulting in a final pixel size of 130 nm in the focal plane, after 2 × 2 binning. For fluid exchange each individual chamber of the ibidi μ-SlideVI^0.5^ were fitted with elbow Leur connector male adaptors (Ibidi) and 0.5 mm silicon tubing (Ibidi). Each imaging acquisition step was performed by adding the corresponding imager strand buffer solution to the sample. Prior to imager exchange, the chamber was washed for 10 min with 1x PBS buffer with 500 mM NaCl. Before the next imager strand buffer solution was added, we monitored with the camera to ensure complete removal of the first imager strand. Sequential imaging and washing steps were repeated for every cell imaged. For each imaging step, 15,000 frames were acquired with 100 ms integration time and a laser power density at the sample of 0.5 kW/cm^2^.

### Super resolution DNA-PAINT image reconstruction

Both P2Y_2_ and αV Images were processed and reconstructed using the Picasso (Schnitzbauer *et al*., 2017) software (Version 0.3.3). The Picasso ‘Localize’ module was used to identify and localise the x,y molecular coordinates of single molecule events from the raw fluorescent DNA-PAINT images. Drift correction and multi-colour data alignment was performed via the Picasso ‘Render’ module, using a combination of fiducial markers and multiple rounds of image sub-stack cross correlation analysis. Localisations with uncertainties greater than 13 nm were removed and no merging was performed for molecules re-appearing in subsequent frames. Super-resolution image rendering was performed by plotting each localization as a Gaussian function with standard deviation equal to its localization precision.

### Protein quantification via qPAINT analysis

To convert the list of *x,y* localisations into a list of *x,y* protein coordinates the data was further processed using a combination of DBSCAN cluster analysis, qPAINT analysis and *k-*means clustering.

First, 21 randomly selected, non-overlapping, 4×4 μm^2^ regions on interest (ROIs) for each type of cell and cell treatment were analysed with a density-based clustering algorithm, known as DBSCAN. To avoid suboptimal clustering results; ROIs were selected such that they do not intersect with cell boundaries and the regions were the same for P2Y_2_ and αV images. Single molecule localisations within each ROIs were grouped into clusters using the DBSCAN modality from PALMsiever (Pengo, Holden and Manley, 2015) in MATLAB (2021a)(Pengo, Holden and Manley, 2015). This clustering algorithm determines clusters based upon two parameters. The first parameter is the minimum number of points (‘minPts’) within a given circle. For minPts we chose a parameter in accordance to the binding frequency of the imager strand and acquisition frame number; in our case this was set to 10 localisations for all the experiments. The second parameter is the radius (epsilon or ‘eps’) of the circle of the cluster of single molecule localisations. This is determined by the localisation precision of the super-resolved images and, according to the nearest neighbour based analysis was ca. to 10 nm for all the images.

For qPAINT analysis we used a custom-written MATLAB (2021a) code. Briefly, localisations corresponding to the same cluster were grouped and their time stamps were used to compile the sequence of dark times per cluster. All the dark times per cluster were pooled and used to obtain a normalised cumulative histogram of the dark times which was then fitted with the exponential function 1 – exp(t/τ_d_) to estimate the mean dark time, τ_d_, per cluster. The qPAINT index (*q*_i_) of each cluster was then calculated as the inverse of the mean dark time, 1/τ_d_.

Calibration was then performed via compilation of all qPAINT indexes obtained from the DNA-PAINT data acquired for each protein type into a single histogram. Only qPAINT indices corresponding to small clusters (i.e., cluster with a maximum point distance of 150 nm) were considered. This histogram was fitted with a multi-peak Gaussian function to determine the qPAINT index for a cluster of single molecule localisations corresponding to one protein (*q*_i1_).

The calibration value obtained with this method was used to estimate the number of P2Y_2_ and αV proteins in all the single molecule localisations clusters identified by DBSCAN, as this corresponds to the ratio between *q*_i,1_ and the qPAINT index of each cluster. Finally, *k*-means clustering was used to recover a likely distribution of the proteins’ positions in each cluster of single molecule localisations, where *k* is equal to the number of proteins in that cluster. This information allowed us to quantify the protein density and level of protein clustering.

### Nearest neighbour analysis

Nearest neighbour distances (NND) for P2Y_2_ - P2Y_2_ and αV-αV were calculated using the recovered P2Y_2_ and αV-protein maps as described above via a custom-written MATLAB (2021a) script. For colocalization analysis, the NND for each protein of one dataset with respect to the reference dataset was calculated (i.e., P2Y_2_ - αV) using a similar MATLAB script. To evaluate the significance of the NND distributions, we randomized the positions of P2Y_2_ and αV for the comparison of P2Y_2_ - P2Y_2_ and αV-αV NND distributions, respectively, and the positions of one of the two proteins for the comparison of the NND between P2Y_2_ - αV protein distributions. The resulting histogram of the nearest neighbour distances for both the experimental data sets and the randomly distributed data was normalized using the total number of NND calculated per ROI to calculate the percentage of the populate with distances smaller than a set threshold value.

### Statistical analysis

For the statistical analysis of number and colocalization of DNA-PAINT images, a minimum of five 4×4 μm^2^ regions obtained from AsPC-1 cells were analysed per condition. For all experiments, normality tests were performed and the non-parametric Kruskal-Wallis test for significance was calculated. All graphs and statistical calculations of experimental data were made using Prism 9.4.1 (GraphPad).

## Acknowledgments

We thank the Barts Pancreatic Tissue Bank (BPTB) for providing pancreatic tissue slides presented in this work. BPTB is supported by Pancreatic Cancer Research Fund and we thank all its members, in particular, Claude Chelala, Christine Hughes, Ahmet Imrali and Amina Hughes for help, as well as Consultant Pathologist Dr Joanne Chin-Aleong and members of Tissue Access Committee and Operations Group. We thank Ann-Marie Baker for her expertise on RNAscope experiments. This work was supported by a Medical Research Council (MRC) iCase award to P.J.M. and R.P.G. from Barts Charity and the MRC Doctoral Training Programme for E.T.B. at Queen Mary University of London (Project MRC0227). N.J.R. acknowledges the QMUL MRC Doctoral Training Program (MR/N014308/1). M.D.J. acknowledges support from the BBSRC (BB/T008709/1) via the London Interdisciplinary Doctoral Programme and S.S. acknowledges financial support from the Royal Society through a Dorothy Hodgkin fellowship (DHF\R1\191019) and a Research Grant (RGS\R2\202038). This work was supported by Cancer Research UK (CRUK) awarded to E.P.C. and R.P.G. (A27781) and a CRUK Centre grant to Barts Cancer Institute (A25137). Human PDAC tumour data were generated by TCGA Research Network (https://www.cancer.gov/tcga) and by the Clinical Proteomic Tumour Analysis Consortium (https://www.proteomics.cancer.gov). The Genotype-Tissue Expression (GTEx) Project was used for the analysis of normal tissue samples (https://gtexportal.org). Diagrams were generated using BioRender.

## Author Contributions

E.T.B., P.J.M. and R.P.G. designed and directed the study; E.T.B. performed and analysed the majority of experiments and bioinformatic analyses; M.D.J. was involved in the imaging and analysis of DNA-PAINT results; Q.W. assisted with 3D sphere assays; E.P.C. generated H2B and Lifeact transduced cells and provided expertise with the 3D invasion assay method; N.J.R. assisted with CRISPR cell line generation; J.G. aided with DNA-PAINT analysis, IF and 3D sphere fluorescent imaging; A.S. provided support with IF staining; H.M.K. provided tissue slides through the Bart Pancreatic Tissue Bank and provided the cell lines used; S.S. provided assistance and expertise with DNA-PAINT experiments; E.T.B., M.D.J., S.S., P.J.M and R.P.G. wrote the manuscript. All authors have read and approved the manuscript.

## Supplemental information

**Supplementary Figure 1.**
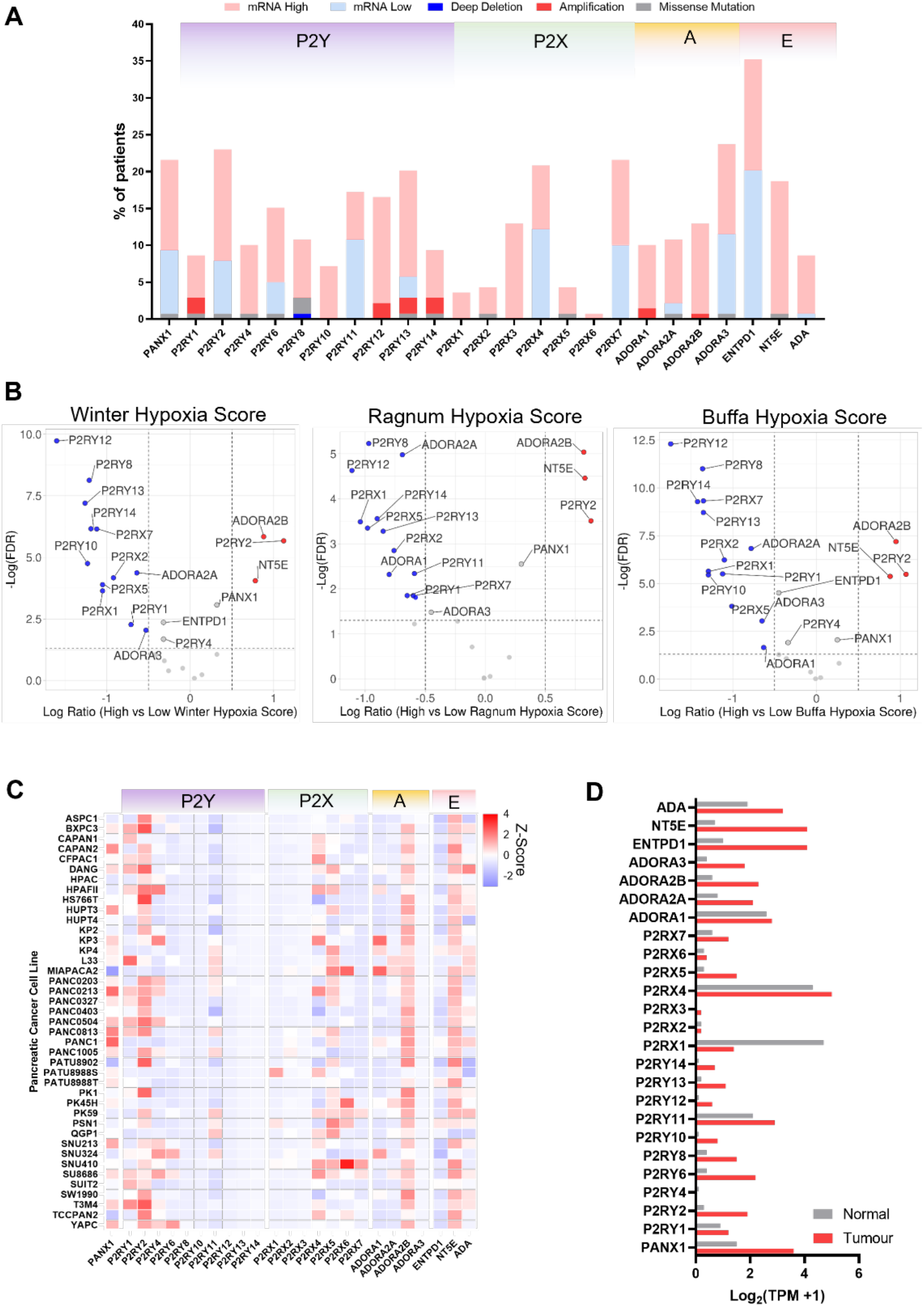
Characterisation of purinergic genes in pancreatic adenocarcinoma. **A** Oncoprint from the PAAD TCGA cohort generated using cBioPortal. mRNA high and mRNA low represent Z-score values of >1 or <-1. **B** Volcano plots for differential expression results of PAAD TCGA patient of high or low hypoxia scores using 3 different hypoxia signatures (Winter, Ragnum and Buffa). **C** Heat map of purinergic mRNA expression data for different PDAC cell lines from CCLE. **D** Comparison of normal versus tumour normalised transcripts per million (TPM) expression of purinergic genes. Data obtained using GEPIA and PAAD TCGA and GTEx.

**Supplementary Figure 2.**
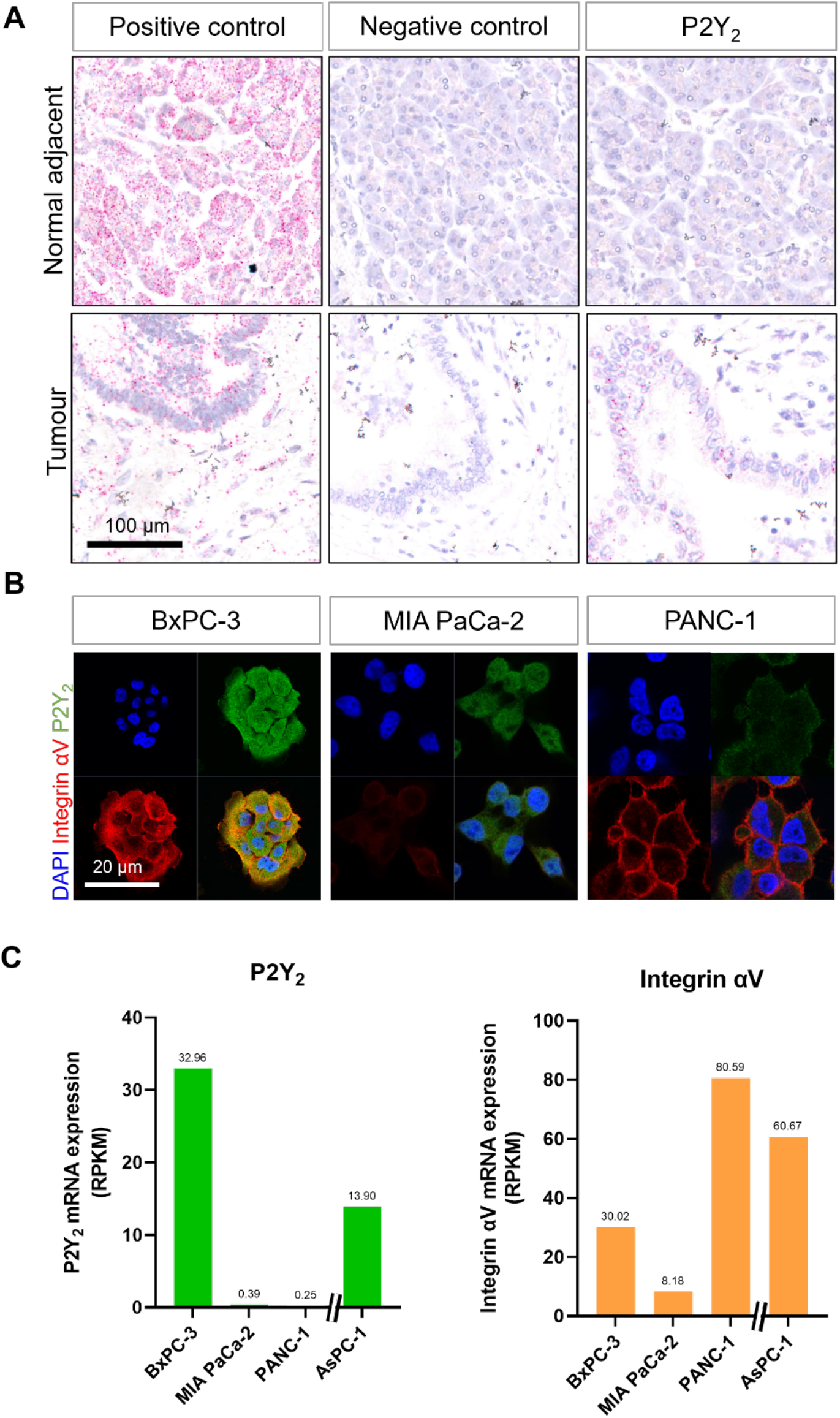
mRNA and protein expression of P2Y_2_ in PDAC cells. **A** RNAscope *in-situ* hybridisation of a positive control (*PIPP*, Cyclophilin B), negative control (*DapB*) and P2Y_2_ mRNA expression in a PDAC tissue slide showing tumour and normal adjacent tissue. **B** IF staining of 3 different PDAC cell lines showing various levels of P2Y_2_ (green) and integrin αV (red) protein expression and the respective reads per kilobase of exon per million reads mapped (RPKM) from CCLE in **C**.

**Supplementary Figure 3.**
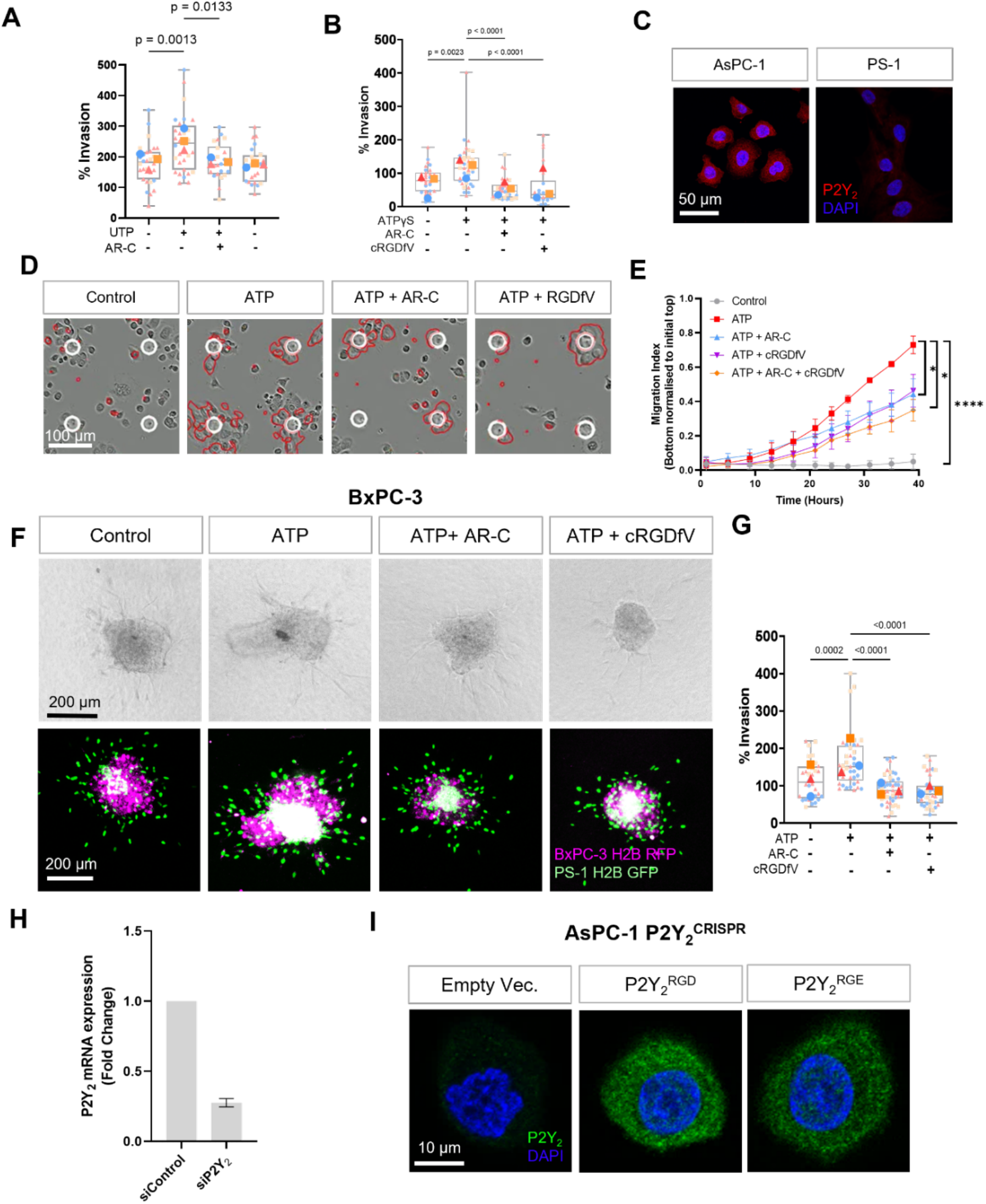
Invasion and migration experiments in PDAC cell lines. **A, B** Quantification of AsPC-1 spheres treated with 100 μM UTP or ATPγS (respectively) in absence or together with 5 μM AR-C or 10 μM cRGDfV. **C** IF staining of P2Y_2_ in AsPC-1 and PS-1 stellate cells. **D** Migration assay with AsPC-1 and 100 μM ATP in absence or together with 5 μM AR-C or/and 10 μM cRGDfV and its quantification (**E**). **F** 3D sphere invasion assay using BxPC-3 cells treated with 100 μM of ATP in absence or together with 5 μM AR-C or/and 10 μM cRGDfV and its quantification (**G**). **H** qPCR of P2Y_2_ expression of siRNA treated cells (control siRNA and P2Y_2_ targeting siRNA) normalised to GAPDH. **I** P2Y_2_ IF staining of AsPC-1 P2Y_2_^CRISPR^ cells transfected with an empty vector, P2Y_2_^RGD^ or P2Y_2_^RGE^ plasmids.

**Supplementary Figure 4.**
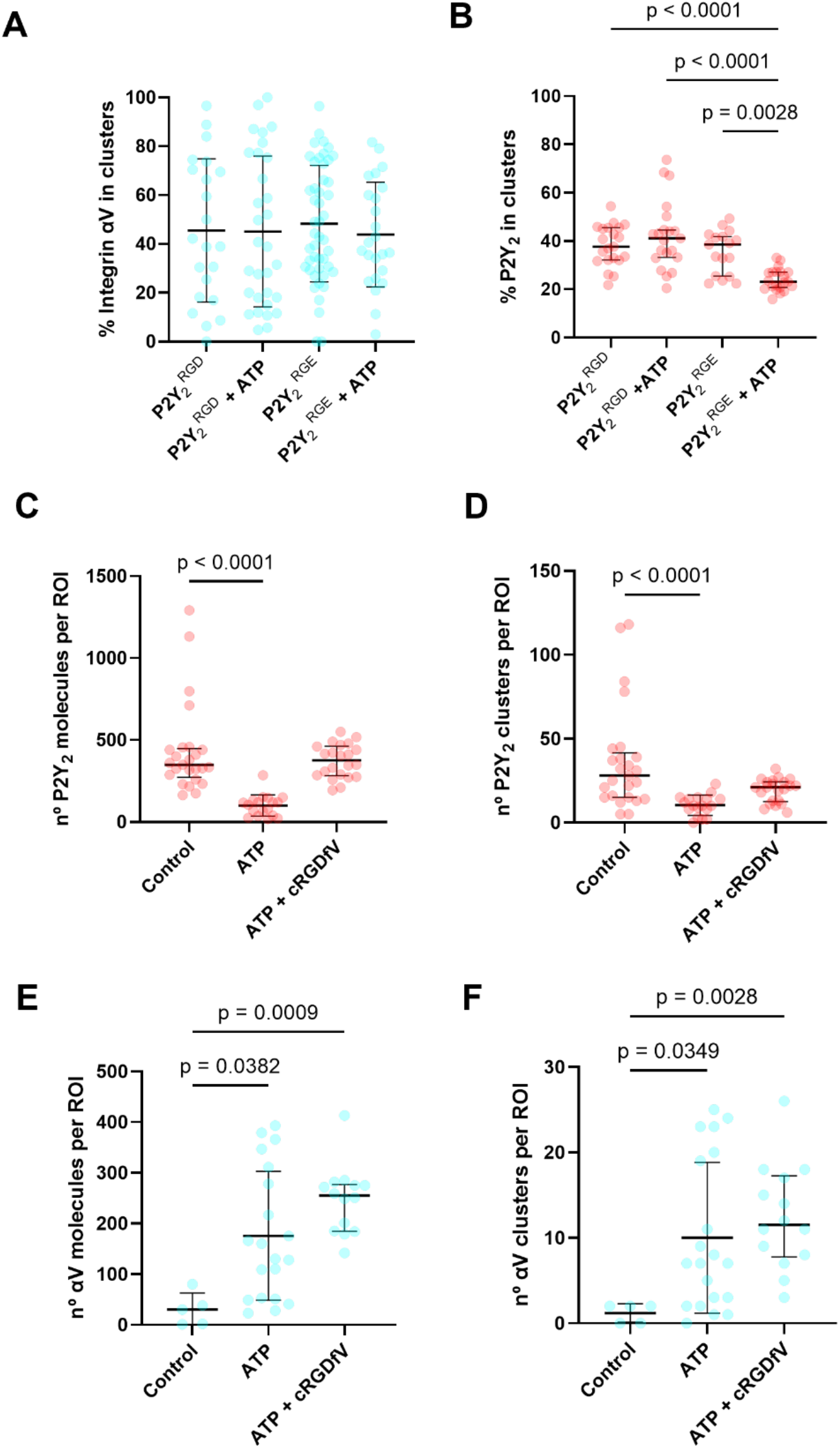
Quantification of P2Y_2_ and integrin αV at the membrane using DNA-PAINT. Percentage of integrin αV (**A**) and P2Y_2_ (**B**) in clusters normalised to the number of proteins (integrin αV or P2Y_2_ proteins respectively) in AsPC-1 P2Y_2_^CRISPR^ cells transfected with P2Y_2_^RGD^ or P2Y_2_^RGE^ and treated with vehicle control or 100 μM of ATP. Non-altered AsPC-1 cells treated with vehicle control or 100 μM of ATP with or without cRGDfV were imaged with DNA-PAINT. The quantification of the number of proteins or protein clusters (>3 proteins) in each region of interest (ROI) are shown in red for P2Y_2_ (**B** and **C** respectively) and in cyan for integrin αV (**D** and **E** respectively).

